# Passive acoustic monitoring of Tasmanian masked owls and swift parrots: an effective tool for conservation actions

**DOI:** 10.1101/2025.08.07.669245

**Authors:** Charley Gros, Matthew H. Webb

**Affiliations:** Bob Brown Foundation, Hobart, Tasmania, Australia; Landscape Recovery Foundation, Hobart, Tasmania, Australia

**Keywords:** passive acoustic monitoring, neural networks, conservation, *Tyto novaehollandiae*, *Lathamus discolor*

## Abstract

The Tasmanian masked owl (*Tyto novaehollandiae* subsp. *castanops*) and swift parrot (*Lathamus discolor*), both endangered species, rely on old forest features that are declining across their ranges in Tasmania, Australia. Their elusive behavior and high mobility make monitoring difficult, hindering conservation actions. Passive acoustic monitoring can greatly increase spatial and temporal survey coverage, though the identification of the species’ vocalisations within large audio datasets remains challenging.

We deployed automated recording units at 108 sites in Tasmania’s native forests to collect a large and representative acoustic dataset. We trained a neural network model to automate call detection for both the Tasmanian masked owl and the swift parrot.

Our model demonstrated high performance, with a sensitivity of 97.7% and specificity of 96.5% for the Tasmanian masked owl, and a sensitivity of 87.5% and specificity of 83.3% for the swift parrot. Through two real-world applications, we illustrated how our method provides detailed quantitative insights into habitat use patterns over extended spatial and temporal scales. This innovative approach enhances both site-specific and population monitoring, enabling more effective and targeted conservation actions for these endangered species.

## 1. Introduction

The Tasmanian masked owl *Tyto novaehollandiae* subsp. *castanops* and swift parrot *Lathamus discolor* are primarily threatened by habitat loss and are high priority for conservation in Tasmania’s native forests (Bell & Mooney, 2002; Owens et al., 2025; Webb et al., 2018). Both species are threatened under the Tasmanian, *Threatened Species Protection Act 1995* and the Australian, *Environment Protection and Biodiversity Conservation Act* 1999. Population and habitat monitoring in remote areas of Tasmania’s native forests is challenging for both the Tasmanian masked owl and swift parrot. Monitoring of swift parrots mainly relies on repeated site visits by observers though detectability is limited due to the species’ high mobility and the interannual spatial variation in breeding habitat (Webb et al., 2012, 2014). Masked owl detection surveys have typically used call broadcasts to elicit a call response from individuals, though this method usually yields low detection rates due to the species’ large territory and infrequent vocalisations (Loyn et al., 2011; Todd et al., 2018). Recently, passive acoustic monitoring has been successfully used for detection of Tasmanian masked owls (Gros et al., 2023) however, the processing of typically large acoustic datasets to identify calls has been tedious and time-consuming.

Passive acoustic monitoring is an emerging alternative to traditional wildlife monitoring surveys (Pérez-Granados & Traba, 2021; Shonfield & Bayne, 2017; Sugai et al., 2019) and typically involves deploying autonomous recording units to passively record sounds on a set time schedule, followed by off-site analysis of the acoustic data. Passive acoustic monitoring can provide a cost-effective and minimally invasive way to sample continuously across multiple locations over extended time periods (Buxton et al., 2018; Darras et al., 2019; Hill et al., 2018; Pérez-Granados & Traba, 2021; Sugai et al., 2019). Acoustic data form a permanent record that can be re-analysed to reduce observer bias (Knight et al., 2017; Sugai et al., 2019), and can be effective for detecting rare species with low detectability (Baroni et al., 2023; Dale et al., 2022; Duchac et al., 2020; Holmes et al., 2014; Pérez-Granados et al., 2018). In avian research, acoustic recorders are used for monitoring population trends (Furnas & Callas, 2015; Wood et al., 2019), studying behaviour (Honig & Schackwitz, 2023; Perrault et al., 2014), and modelling habitat associations (Campos-Cerqueira & Aide, 2016; Metcalf et al., 2022). Integrating traditional survey methods with acoustic monitoring has the potential to be a valuable tool for overcoming challenges in quantifying habitat use and breeding success for cryptic and highly mobile threatened species like the Tasmanian masked owl and swift parrot.

The time-consuming nature of manual species identification using passive acoustic monitoring collected across extensive spatial areas and typically involving hundreds or thousands of hours of recordings has driven the need for streamlined acoustic data processing and reliable methods to automatically identify species. Several approaches have been proposed worldwide for automatic bird call detection, including decision trees (Digby et al., 2013), spectrogram cross-correlation (Arvind et al., 2023; Katz et al., 2016), hidden Markov models (Wildlife Acoustics, Inc., 2021), and, most recently, deep learning using neural networks (Nolan et al., 2022; Ruff et al., 2021; Zhong et al., 2020). Neural networks typically represent data as 2D spectrograms and frame the problem as an image classification task. Neural networks like BirdNET (Kahl et al., 2021) are increasingly used in avian research and citizen science projects (Clark et al., 2023; Wood et al., 2023). The strength of neural networks lies in their ability to learn hierarchical representations, enabling them to identify relevant features without relying on predefined settings. Early layers learn basic patterns, while deeper layers capture abstract shapes that are crucial for distinguishing bird species and are robust to variations in spectrograms. However, training neural networks on highly imbalanced datasets, especially for rare or elusive species, remains a key challenge.

We developed a passive acoustic monitoring pipeline for Tasmanian masked owl and swift parrot calls, including: (i) data collection from a broad and representative range of rainforest and native eucalypt forest across Tasmania, (ii) automated identification of target calls in field recordings, and (iii) data analysis through two real-life case studies. We assessed the robustness of the method using an independent, multi-site dataset with a range of vegetation types, seasonal variations, and non-target species. Our study highlights how this approach can enhance our understanding of the ecology of both the Tasmanian masked owl and swift parrot and contribute to the development of more effective conservation actions.

## 2. Methods

In this section, we outline the process of training a model to automatically detect the calls of the Tasmanian masked owl and swift parrot, covering data collection, preprocessing, model training, and validation. We also assess the performance of this approach through two practical applications for monitoring these species.

### 2.1. Data collection

We deployed passive acoustic recorders in native forests distributed across Tasmania, covering a range of forest types and elevations between June 2021 and February 2024 (Figure 1). Most recordings (>95 %) were obtained using Song Meter Mini recorders (Wildlife Acoustics, Maynard, Massachusetts). Other recording equipment included BAR-LT (Frontiers Lab, Brisbane, Australia) and Chorus (Titley, Brendale, Australia) recorders. Acoustic data were stored as hour-long WAV files with a sampling rate of 22,050 Hz.

**Figure 1.**
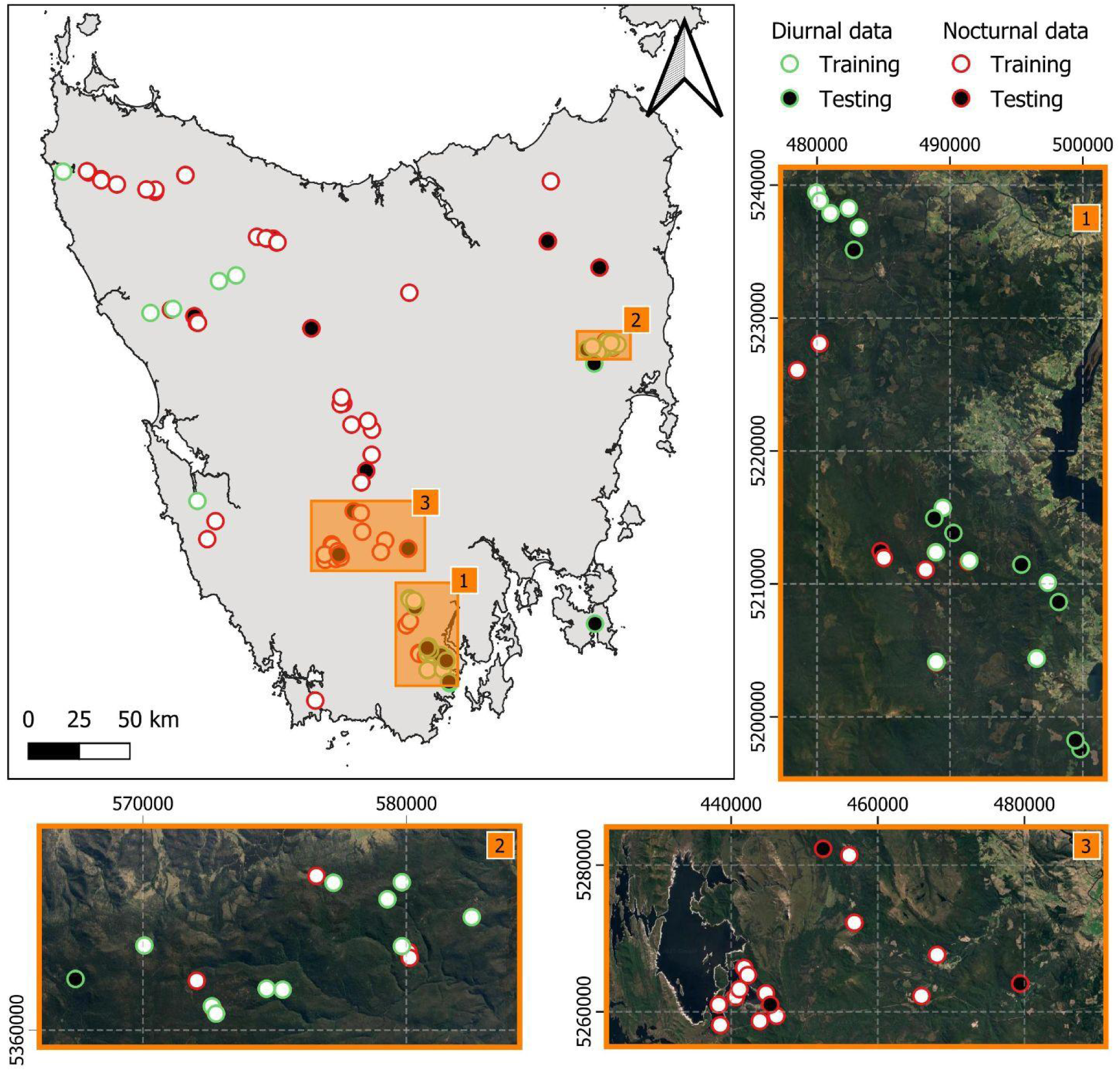
Data spatial distribution across Tasmania. Each dot represents a survey site where acoustic recorders were deployed to collect diurnal data (circled in green) and to collect nocturnal data (circled in red). White dots show locations where the data collected were used to train and optimise the models. Black dots show locations where data were collected independent of the training dataset (i.e., at least 500 m away), and were used to evaluate the models.

Diurnal data were collected for swift parrot monitoring during their breeding season (October to February), with variable time recording schedules depending on the location and forest type (n_site_ = 41; Figure 1). Nocturnal data, collected for Tasmanian masked owl monitoring, followed a consistent time schedule from sunset to sunrise (n_site_ = 67; Figure 1). The target species were not detected at all sites, but sites with no detection were valuable for training data, expanding the “negative” class to include vocalisations of other species and the local soundscape.

We randomly selected ten sites for the nocturnal data and thirteen sites for the diurnal data to be used for an independent evaluation of the models, while the remaining data were used to compile a training dataset (Figure 1).

We focused on automatically detecting the screech call of the Tasmanian masked owl, and three distinctive swift parrot calls including: the adult ‘flight’ call, the adult ‘warble’ call, and the ‘chick begging’ call. Figure 2 shows spectrograms for each of the four target calls. Todd et al. (2018) described the adult Tasmanian masked owl screech call as a mean call duration of 1.6 s and a median frequency of 1,784 Hz. The swift parrot flight call is a loud, high-pitched, sharp, metallic ‘chit’, repeated rapidly. The warble call, heard during feeding or social interactions, features complex, cascading notes with rising and falling pitch. The swift parrot chick begging call is a repetitive ‘cheep’ similar to a typical parrot begging call.

**Figure 2.**
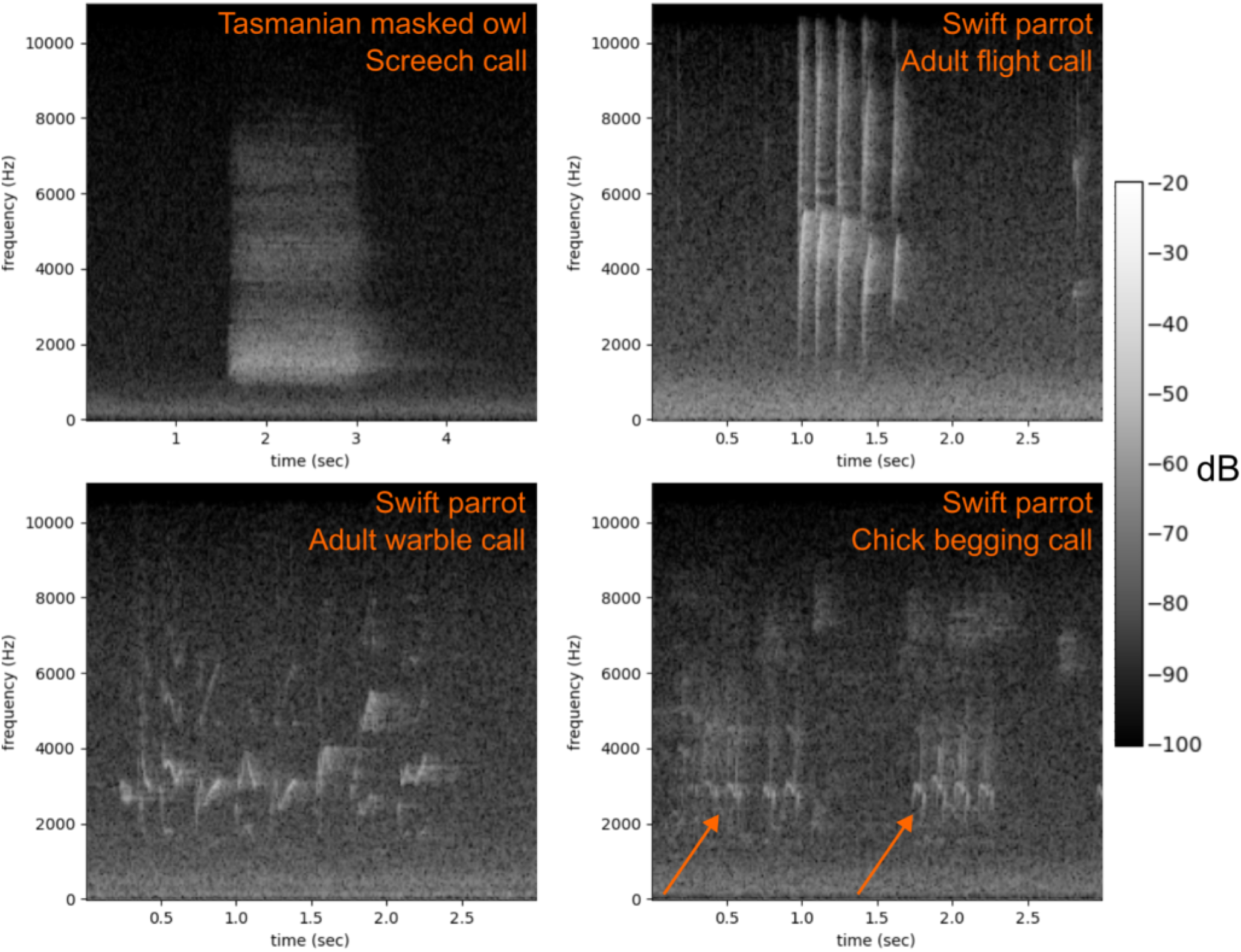
Example spectrograms of Tasmanian masked owl and swift parrot calls.

### 2.2. Model generation and evaluation

In this section, we provide a step-by-step account of how we trained, iteratively improved, and evaluated a neural network model to automatically detect the calls of the two target species.

#### 2.2.1. Training data compilation

The training data comprises short audio clips of the target calls, along with “negative” sounds (e.g., non-target species, background noise). Training data was “weakly labelled”, i.e. we provided the correct label for each audio clip but not the delineation of the vocalisation within the spectrogram. The training data compilation was an iterative process. For the swift parrot, we compiled an initial training dataset by manually extracting audio clips containing swift parrot calls from raw data collected at locations where the species was known to be present. For the Tasmanian masked owl, we used Kaleidoscope Lite software (v5.4.8) (Wildlife Acoustics, Inc., 2021) to generate an initial set of the target calls. Kaleidoscope allowed us to detect sounds that meet some criteria, i.e. sound characteristics. We developed these criteria empirically to maximise the detection of Tasmanian masked owl screech calls (i.e., 0.6 – 2s in duration, 1–2kHz in frequency, with a maximum inter-syllable gap of 0.1s) (Gros et al., 2023). We reviewed the audio clips pre-selected by Kaleidoscope and compiled “positive” and “negative” samples for the Tasmanian masked owl screech call. To increase the diversity of the “negative” samples, we also extracted audio clips at a random offset within randomly selected raw field recordings. We reviewed each randomly selected audio clip to ensure that no “positive” sample was misclassified. This initial training data was used to train a first model for each species. This automated recogniser was used on raw data subsequently collected, which increased the size of the training dataset. The final training dataset used for this study contains: (i) 4,030 Tasmanian masked owl screech calls and 5,800 negative samples for the nocturnal data, and (ii) 4,602 swift parrot adult flight calls, 1,236 adult warble calls, 5,734 chick begging calls, and 10,002 negative samples for the diurnal data.

#### 2.2.2. Preprocessing

The audio clips were trimmed to 5-second duration for the nocturnal model and 3-second duration for the diurnal model to be sufficiently long to encompass target calls, yet short enough to deliver high temporal accuracy during predictions. Following previous approaches, e.g. (Kahl et al., 2021; Ruff et al., 2021; Zhong et al., 2020), the model’s input is a spectrogram that is treated as a monochrome image, which means that the model is trained to tackle an image classification task. Spectrograms were generated using a Hann window with a window length of 2048, 50% window overlap, and 512 audio samples per spectrogram window. All spectrograms covered a frequency range from 0 to 11.025 kHz, with the upper frequency limit reflecting the Nyquist frequency of the 22.05 kHz sample rate. Spectrograms were saved as grayscale images, 257 pixels x 429 pixels.

#### 2.2.3. Model training

We used a model architecture with an 18-layer residual network architecture, or ResNet-18, initially proposed by He et al. (2015). ResNets have produced good classification performance on various passive acoustic monitoring classification tasks (e.g., Allen et al., 2021; Batist et al., 2024; Bergler et al., 2022; Dufourq et al., 2022).

The models were initialised to weights pre-trained on an extensive image dataset called ImageNet (Deng et al., 2009) which is designed for generic image classification tasks. This pre-trained model was then re-purposed for our specific task by fine-tuning the model’s weights on our training datasets. This approach is effective because it prevents having to train the whole model with random initialisation of all parameters (n ∼ 11 million), which would typically require a much larger training dataset than currently available (Chollet, 2021; LeCun et al., 2015). Data augmentation is commonly used when training deep learning models to mitigate the risk of over-fitting (Chollet, 2021). Incorporating domain-specific data augmentation is crucial to accommodate unforeseen variations in real-world samples. During the models’ training, three data augmentation operations were performed: (i) drawing horizontal and vertical bars over the spectrogram and replacing with the mean image value, to generate random occlusions; (ii) adding gaussian noise, to generate random background noise; and (iii) adding randomly selected gain level, to simulate varying sound intensities.

The models were trained for 50 epochs with a random 5:1 training-validation split and a batch size of 64. Loss was calculated with the categorical cross-entropy function. To train the models, the stochastic gradient descent optimisation algorithm (Bottou, 2010) was used with an initial learning rate of 10^-2^. The learning rates were multiplied by 0.1 once every five epochs in order to decrease the learning rate over the course of the training, i.e. weights fine-tuning. Early stopping with a cooldown of two epochs prevented over-fitting. The above-mentioned parameters were selected through a non-exhaustive set of preliminary experiments that compared the validation metrics curves, e.g., model’s precision on the validation dataset. Our overall model training process largely followed current best practices in deep learning, which are well detailed in Chollet (2021).

The training pipeline was implemented in Python 3.10 and included the following libraries: “PyTorch” (v2.1.0) (Paszke et al., 2019) for the neural network component, and “opensoundscape” (v0.10) (Lapp et al., 2023) for audio processing. The model training was carried out on a single NVIDIA Tesla P100 GPU with 16 GB RAM memory and took approximately eight and 14 hours, for the Tasmanian masked owl and the swift parrot models, respectively.

#### 2.2.4. Model inference

At inference time, each raw field recording was loaded, re-sampled to 22.05 kHz if applicable, and split into three second clips for the diurnal data and five seconds for the nocturnal data. Acoustic data was then converted to a spectrogram and passed through our model, which yields prediction probabilities for each modelled class. Optimal thresholds for converting detection probabilities into binary classifications (e.g., swift parrot chick begging call) were those that maximise the Area Under the Curve of Receiver Operating Characteristics (AUC-ROC) in the validation dataset. Contrary to the model’s training which requires high computational power such as that offered by a GPU, model prediction can run on a standard consumer-like laptop.

#### 2.2.5. Model evaluation

To assess the models’ performance, a testing dataset was manually annotated: 100 h (n_site_ = 10) and 20 h (n_site_ = 13) of raw field recordings for the Tasmanian masked owl and the swift parrot models, respectively. This dataset is independent from the dataset used for the model’s training, i.e. the data were collected at different sites (see Figure 1). For each class, two metrics were computed: the sensitivity and the specificity across all one-hour files in the testing dataset. Sensitivity measures the model’s ability to detect true positives, while specificity indicates its ability to avoid false positives (see Appendix A for details). To assess the impact of newly collected training data on model performance, the performance of our final models were compared to those previously trained using the same approach but with less data. We also compared the performance of the Tasmanian masked owl model with the approach we used in a previous study by Gros et al. (2023) employing the Kaleidoscope Lite software (v5.4.8) (Wildlife Acoustics, Inc., 2021). Additionally, given BirdNET’s growing use in identifying bird vocalisations in acoustic data (Kahl et al., 2021), we compared our models to BirdNET outputs (v2.4 with default parameters) for the two target species.

### 2.3. Applications

We applied our trained model in two real-life case studies to evaluate the effectiveness of our passive acoustic monitoring pipeline.

#### 2.3.1. Tasmanian masked owl

In this case study, we aimed at characterising the spatial and temporal distribution of Tasmanian masked owl calls within and around a logging site to better understand their habitat use. The Tasmanian masked owl was detected in a logging compartment in the Florentine Valley in southern Tasmania in 2022 using a single acoustic recorder. To further explore the nature of the detection, we deployed an array of 18 acoustic recorders within and around the logging site in early 2024, recording from sunset to sunrise. The acoustic recorders were arranged in a spatial configuration following a hexagonal grid with a distance between recorders of about 300 m. The recorders were placed in rainforest, mixed forest and wet eucalyptus forest with recent disturbances including wildfire and/or logging.

#### 2.3.2. Swift parrot

In this case study, we aimed at quantifying swift parrot vocal activity within a logging site, and potentially identifying breeding and breeding success. In December 2023, swift parrots were observed feeding on abundant Blue gum *Eucalyptus globulus* flowering within a logging compartment in Tasmanian Southern forests. The area comprised wet eucalypt forest dominated by Stringybark *E. obliqua*, Mountain ash *E. regnan* and Blue gum. Suitable nesting hollows for swift parrots were present and swift parrot nesting behaviour was observed within the logging site. An acoustic recorder was placed near the base of a potential nesting tree on 13 December 2023, scheduled to record from sunrise to sunset and was retrieved on 29 January 2024.

## 3. Results

### 3.1. Model evaluation

We assessed overfitting risk by computing evaluation metrics throughout model training. During the training, the curves on the validation dataset closely followed those on the training dataset (see Appendix B), indicating consistency. Final models achieved high AUC-ROC scores on the validation dataset i.e., 0.99 for Tasmanian masked owl screeches, 0.99 for swift parrot adult flight calls, 0.96 for adult warble calls, and 0.98 for chick begging calls (see Appendix C). ROC curves on the validation dataset illustrate excellent model fitting performance.

We evaluated the trained models on an independent dataset, separate from the training and validation sets. The Tasmanian masked owl model achieved excellent results, with a sensitivity of 97.7% and a specificity of 96.5%. Table 1 compares our model (named “Ours, n_train+_=4,030”) to the v2.4 BirdNET software (Kahl et al., 2021), to the unsupervised method Kaleidoscope Lite used in a previous study (Gros et al., 2023), as well as to our own approach with fewer training samples, labelled “Ours, n_train+_=1,788”. Our new model outperformed previous approaches, which exhibited higher numbers of false positive and negative detections. For instance, Kaleidoscope Lite showed a sensitivity of 74.4% and specificity of 84.2%. The BirdNET model failed to reliably detect the Tasmanian masked owl, detecting only 14.0% of instances when the species was vocalising in the testing dataset.

**Table 1:** Evaluation metrics on the independent testing dataset. The sensitivity and specificity for the Tasmanian masked owl were calculated for the outputs of Kaleidoscope Lite and BirdNET, as well as for our approach trained with different numbers of positive samples (n=1,788 vs. 4,030). For the swift parrot, these metrics were computed for BirdNET and our approach trained with varying numbers of positive samples (n=4,002 vs. 8,110 vs. 11,572). The optimal value for both sensitivity and specificity is 100.

| Species | Call | Model | Sensitivity | Specificity |
| --- | --- | --- | --- | --- |
| Tas. masked owl | Screech | Kaleidoscope Lite | 74.4 | 84.2 |
|  |  | BirdNET v2.4 | 14. | 100. |
| | | Ours, $n_{\text{train}+}=1,788$ | 93.0 | 77.2 |
| | | Ours, $n_{\text{train}+}=4,030$ | 97.7 | 96.5 |
| Swift parrot | Flight | Ours, $n_{\text{train}+}=4,002$ | 90. | 100. |
| | | Ours, $n_{\text{train}+}=8,110$ | 90. | 100. |
| | | Ours, $n_{\text{train}+}=11,572$ | 90. | 100. |
| | Warble | Ours, $n_{\text{train}+}=4,002$ | 71.4 | 83.3 |
| | | Ours, $n_{\text{train}+}=8,110$ | 64.3 | 83.3 |
| | | Ours, $n_{\text{train}+}=11,572$ | 71.4 | 83.3 |
| | Chick | Ours, $n_{\text{train}+}=4,002$ | 100. | 0. |
| | | Ours, $n_{\text{train}+}=8,110$ | 92.9 | 16.7 |
| | | Ours, $n_{\text{train}+}=11,572$ | 100. | 83.3 |
|  | All | BirdNET v2.4 | 31.3 | 100. |
| | | Ours, $n_{\text{train}+}=4,002$ | 87.5 | 50. |
| | | Ours, $n_{\text{train}+}=8,110$ | 83.3 | 50. |
| | | Ours, $n_{\text{train}+}=11,572$ | 87.5 | 83.3 |

For the swift parrot, increasing the number of training samples did not clearly improve the model’s performance in detecting the flight call and warble call (see Table 1). However, it notably enhanced the model’s ability to avoid false positives, with specificity values of 0%, 16.7%, and 83.3% for the models trained with 4,002, 8,110, and 11,572 positive samples, respectively. Overall, our latest swift parrot model showed better performance detecting the flight call (sensitivity of 90.0% and specificity of 100.0%) and chick begging call (100.0% and 83.3%), compared to the warble call (71.4% and 83.3%). While BirdNet achieved perfect specificity (100. %) with no false positives, it missed many swift parrot calls (i.e., high number of false negatives), resulting in a low sensitivity of 31.3%.

### 3.2. Applications

For the case studies, we collected 614 hours of nocturnal data from the Florentine Valley site, and 501 hours of diurnal data from the Southern forests site. The data processing took 15.5 hours for the nocturnal dataset (i.e., about 40 s per one-hour of field recording), and 8.8 hours for the diurnal dataset (i.e., 60 s per one-hour of field recording) on a consumer-like laptop. The processing steps include: (i) loading the acoustic data, (ii) generating the spectrogram images, (iii) classifying the images with the trained model, and (iv) writing the output, i.e., an audio clip and its spectrogram for each positive detection.

The models yielded 875 apparent detections for the Tasmanian masked owl, and 5,675 for the swift parrot. We reviewed the apparent detections by visually inspecting the spectrograms and listening to the audio clips, which took 2.2 hours for the Tasmanian masked owl, and 8.5 hours for the swift parrot. This review allowed us to confirm that 95.2% of the Tasmanian masked owl detections and 88.4% of the swift parrot detections were correct.

#### 3.2.1. Tasmanian masked owl

The recorders at the Florentine Valley site were collected after 23 nights of data collection. However, some recorders’ batteries ran out before the end of the intended recording period, resulting in a slight variation in the number of nights recorded across the devices, ranging from 18 to 23 nights. The Tasmanian masked owl was detected at every site at least once during the survey period. The number of calls detected varied across the survey area, from two calls at site 16 to 177 calls at site 9. To measure how consistently the species was detected, we calculated the nightly detection rate at each site (defined as the proportion of nights with detections divided by the total number of recorded nights). The nightly detection rate also differed widely between sites, ranging from 10.5% at sites 2 and 16 to 95.7% at site 9. Figure 3 shows the spatial variation in vocal activity across the survey area. Overall, vocal activity was most frequent and consistent at sites 9 and 13, while only a few detections were recorded at sites 2, 6 and 16. Appendix D provides a description of the habitat types at each site and a summary of the survey results.

**Figure 3.**
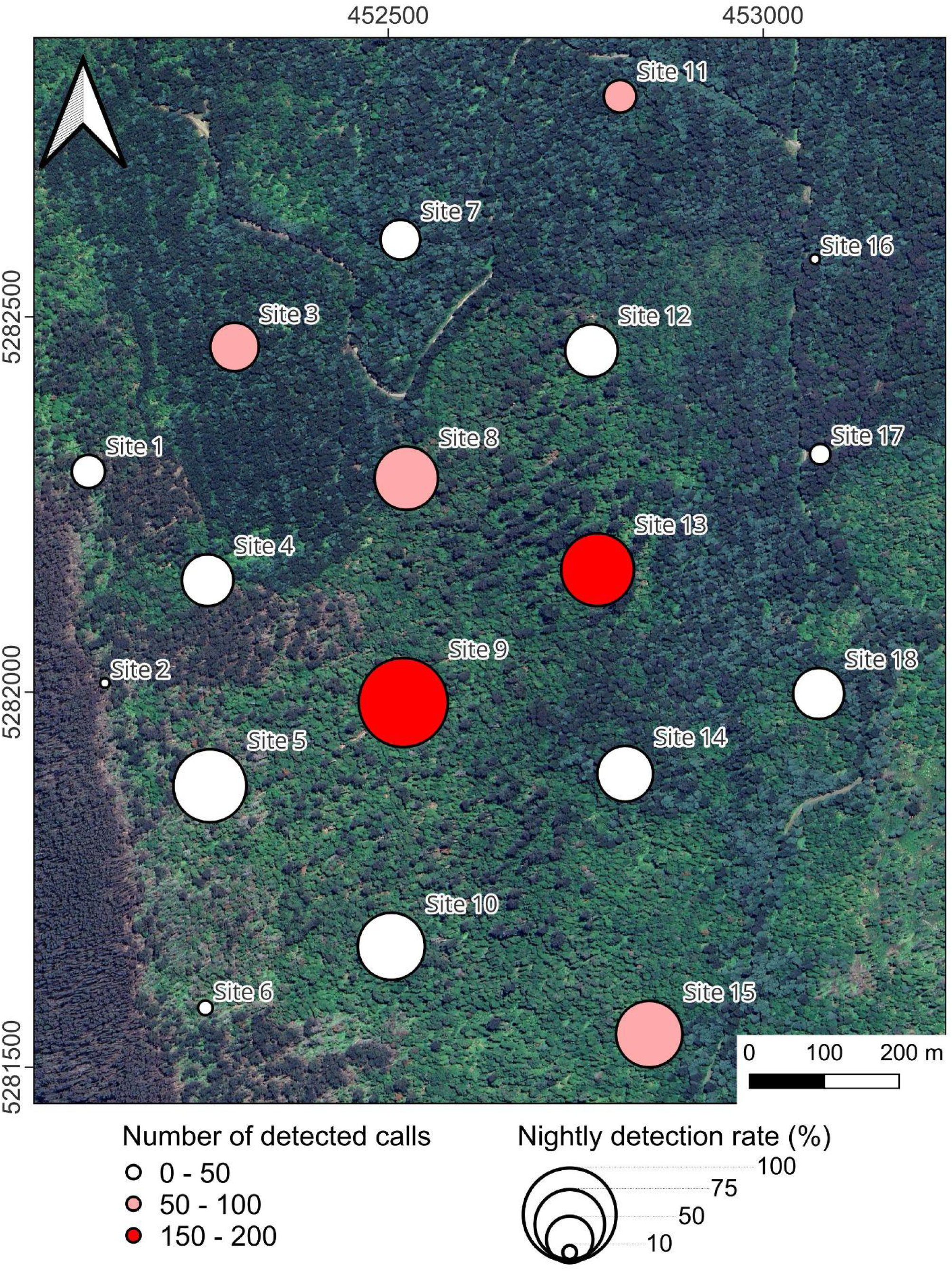
Spatial distribution of Tasmanian masked owl calls at the Florentine Valley site. The vocal activity is measured at each site by the number of calls (from white to red), and by the nightly detection rate (size of the circle, up to 100 %).

The temporal distribution of Tasmanian masked owl calls also varied across sites (Figure 4). For example, detected calls spanned nocturnal hours at sites 18 and 9. In contrast, the majority of calls detected at site 15 were near dusk and dawn. Detections were sparse at some sites (e.g., site 18 on Figure 4), while extremely consistent at others (e.g., site 9). Similarly, the Tasmanian masked owl was detected at dusk and/or dawn throughout the survey period at site 15, with the exception of 8 to 13 March.

**Figure 4.**
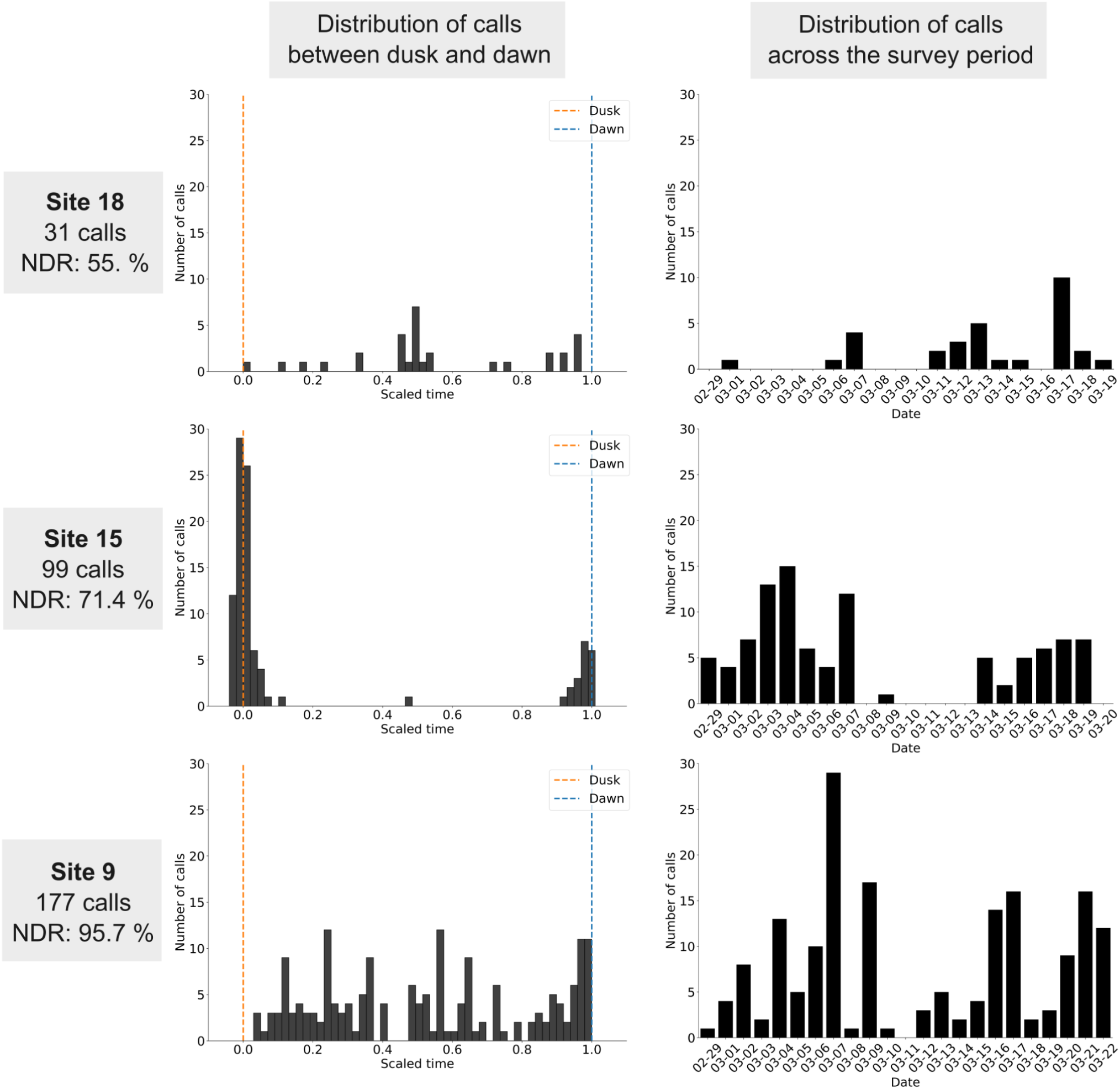
Temporal distribution of Tasmanian masked owl calls at three survey sites. The number of calls is represented for three sites (each row) in two ways: (i) between dusk and dawn (left), and (ii) per night across the survey period (right). Abbreviation: NDR, nightly direction rate.

#### 3.2.2. Swift parrot

Adult swift parrot flight and warbling calls were consistently detected between 14 December 2023 and 15 January 2024, with calls peaking in frequency on 2 January (Figure 5). Adult flight calls were six times more frequent than warble calls. Chick begging calls were detected between 19 December and 13 January, indicating the presence of nesting swift parrots. Prior to 2 January, these calls were rare, with only a few detected each day. Between 2 and 10 January, chick begging calls were frequently detected throughout the day. Adult calls were rarely detected after the last chick begging call on 13 January.

**Figure 5.**
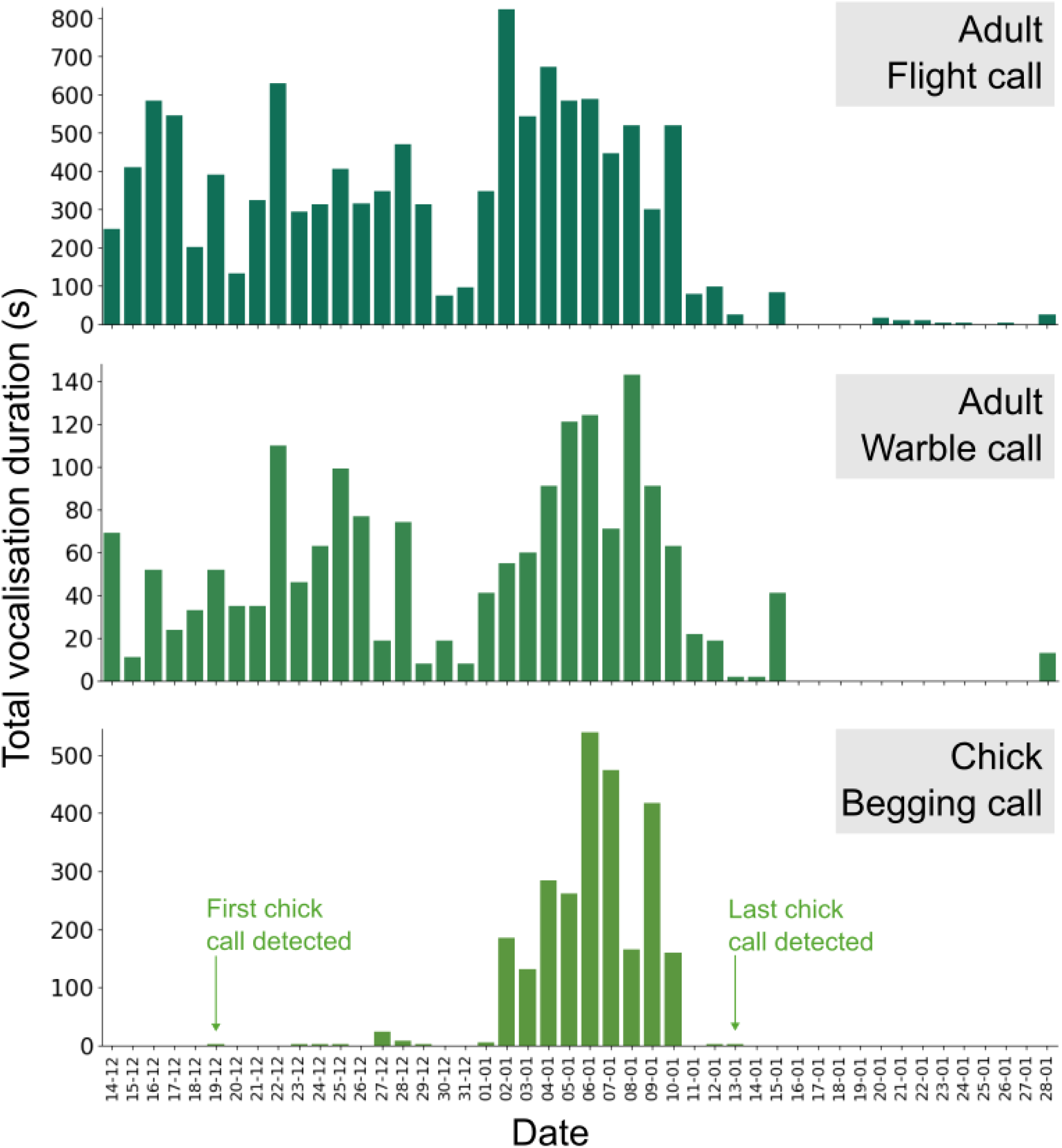
Distribution of swift parrot calls at the nest tree in the Southern forests logging site. For each type of swift parrot call, a bar plot represents the total duration of calls per day that were detected by our model.

## 4. Discussion

Integrating our neural network models into a passive acoustic monitoring program for the swift parrot and the Tasmanian masked owl is transforming our understanding of their distribution and ecology. The strong performance of our models allows efficient analysis of large datasets across broad timeframes and landscapes which was not previously feasible. Our approach has the potential to better inform land management decisions, particularly for activities that involve land clearing or habitat degradation such as forestry or mining. Our method is readily transferable to other rare and/or elusive fauna species that can be detected by distinctive calls and where data deficiencies limit conservation actions.

The potential of passive acoustic recorders for conservation management has been hampered by the availability of efficient automated detection methods to process large datasets. Our study shows that neural networks can streamline the processing of acoustic data, enabling long-term, large-scale monitoring of the Tasmanian masked owl and swift parrot. The heterogeneity of the training dataset, incorporating both regional and seasonal variability, is crucial for capturing the diversity of target and non-target calls. Our models’ performance improved as the number of training samples increased, particularly for the swift parrot chick begging call, the most complex target call in our study. Preliminary experiments revealed that data augmentation (random spectrogram occlusions, background noise variations, and random gain adjustments) significantly enhanced our model training. Future training could explore additional augmentation techniques, such as time-stretching, pitch shifting, and audio mixing (Dufourq et al., 2021; Moummad et al., 2024; Zhong et al., 2020).

Validation on an independent dataset showed our model outperformed Kaleidoscope Lite (Gros et al., 2023), which produced frequent false positives, mainly from brushtail possums (*Trichosurus vulpecula*), whose calls closely resemble the Tasmanian masked owl screech call. BirdNet is an open-source software for automated recognition and acoustic data processing (Kahl et al., 2021), and is increasingly used in passive acoustic monitoring research (e.g., Bota et al., 2023; Pérez-Granados, 2023; Wood et al., 2023). Covering over 6,000 species globally (v2.4, June 2023), it can be a powerful tool for broad-scale biodiversity assessments. In our testing dataset, BirdNet produced almost no false positives, but it frequently failed to detect the swift parrot and Tasmanian masked owl when present, a critical limitation when monitoring rare species. In contrast, our model distinguished between swift parrot call types (i.e., adult flight calls, warble calls, and chick begging calls), providing valuable insights into habitat use and breeding.

Traditional survey methods for the Tasmanian masked owl, such as call-playback (Todd et al., 2018) and radio-tracking (Young et al., 2021), are generally resource intensive, are limited to small sample sizes and small study areas, or contain inherent uncertainties relating to the probability of detection. Our model offers a platform to process and analyse large datasets, allowing us to investigate the species’ occupancy, ecology and distribution, which have hindered the application of passive acoustic monitoring at large spatial scales. We demonstrated the ability to effectively quantify and characterise Tasmanian masked owl vocal activity across multiple spatial scales, providing insights into the species presence in landscapes or fine-scale habitat use. For example, in our case study, the distribution of calls at Site 15 around dusk and dawn suggests the species was nesting and / or roosting nearby. Similarly, insights into the species’ habitat use can be inferred when comparing the high and consistent vocal activity at Site 9 and Site 13 (tall eucalypt trees with an open rainforest understory) with the very limited number of detections at Site 2 (recently affected by wildfires) and Site 16 (young eucalypt regrowth following recent clearfell logging).

Extensive habitat loss within the swift parrot breeding range is considered the primary threat to the species’ population (DCCEEW, 2024), and habitat loss is ongoing (Owens et al., 2025; Webb et al., 2018). Quantifying the species’ dynamic spatial distribution and breeding success in Tasmania is considered essential for conservation planning and informed management actions. The swift parrot case study demonstrates how passive acoustic monitoring can provide essential and detailed insights into the species’ breeding distribution, habitat use and breeding success. Our approach using passive acoustic monitoring can provide near real-time data on swift parrot presence and breeding activity in Tasmania’s native eucalypt forests including in active logging and land clearing operations.

The application of our neural network models to passive acoustic monitoring are yielding new insights into the conservation ecology of the Tasmanian masked owl and the swift parrot. For the Tasmanian masked owl, the analysis of acoustic data is assisting with understanding broad-scale distribution and habitat use by this species, and enabling the identification of fine-scale critical habitat features such as nesting and roosting hollows. For the swift parrot, our approach offers considerable insights into swift parrot feeding site use, nesting locations, nesting attendance, and breeding outcomes, as well as how habitat availability and configuration influence these factors which may be critical for effective conservation. Integrating our approach with traditional survey methodologies is likely to further increase the value of information collected and assist in contextualising data collected from other survey techniques. The metrics quantified in our study are directly applicable to decision-making processes and management actions at local scales such as logging sites, and informing conservation policy, planning and strategies. The integration of passive acoustic monitoring and neural network models enables species monitoring at spatial and temporal resolutions previously unattainable, offering a scalable framework with immediate application to other at-risk and data-deficient species in Tasmania, such as Blue-winged parrot *Neophema chrysostoma* and Tasmanian azure kingfisher *Ceyx azureus* subsp. *diemenensis*, addressing long-standing research limitations and a valuable step forward for their conservation.

## Appendices

## Appendix A: Evaluation metrics of model’s performance

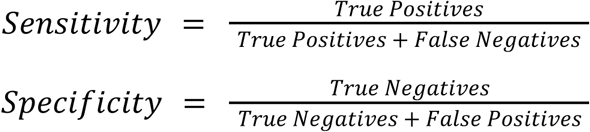

## Appendix B: Model training charts.

At each training step, the model was evaluated on both the training and validation datasets, and the F1-score metric computed. The optimal F1-score value is 1

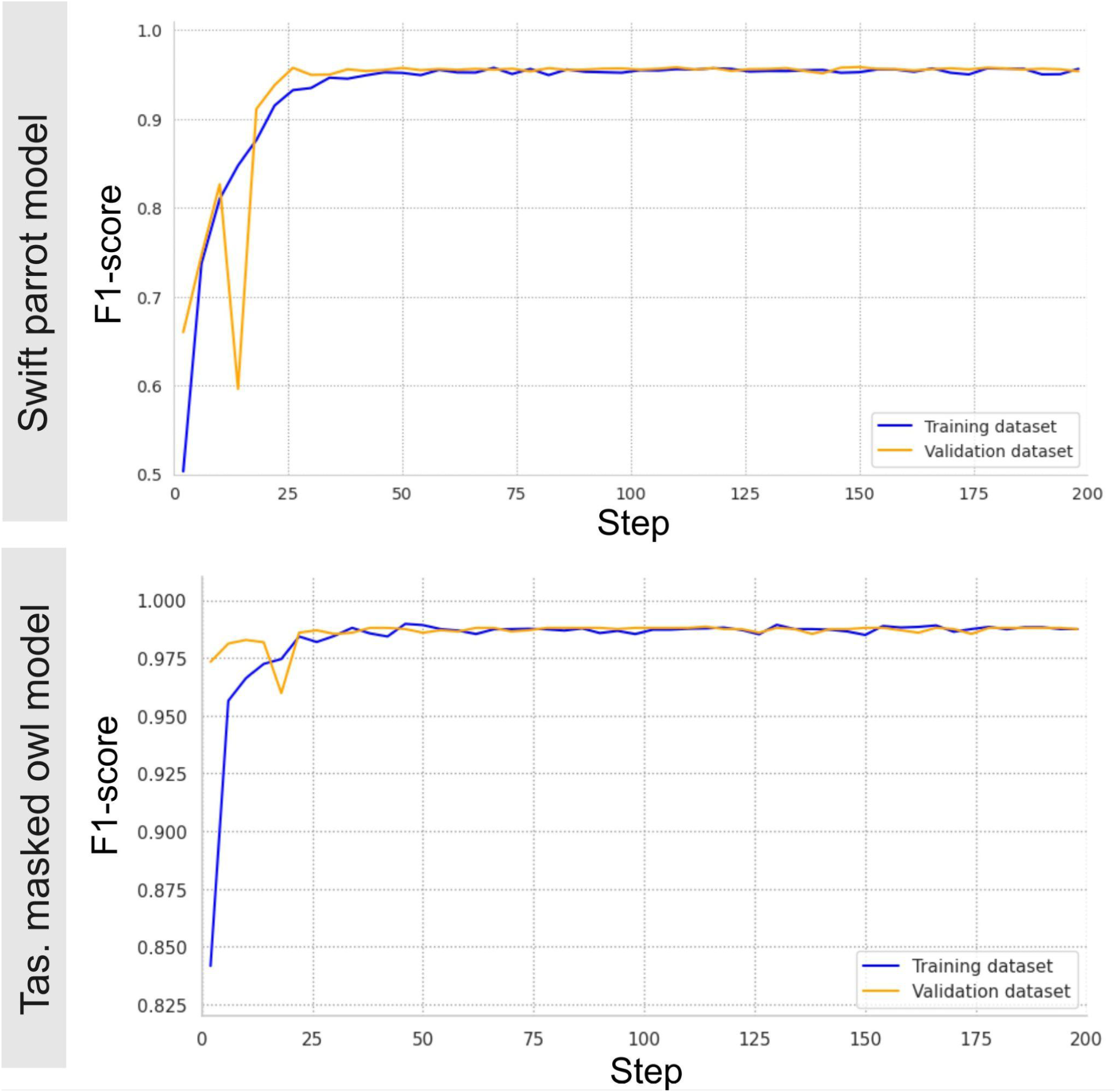

## Appendix C: Receiver operating characteristic curves on the validation dataset.

Each curve is associated with one of the four target calls. The Area Under the Curve (AUC) and the optimal (Opt.) threshold is also indicated for each call

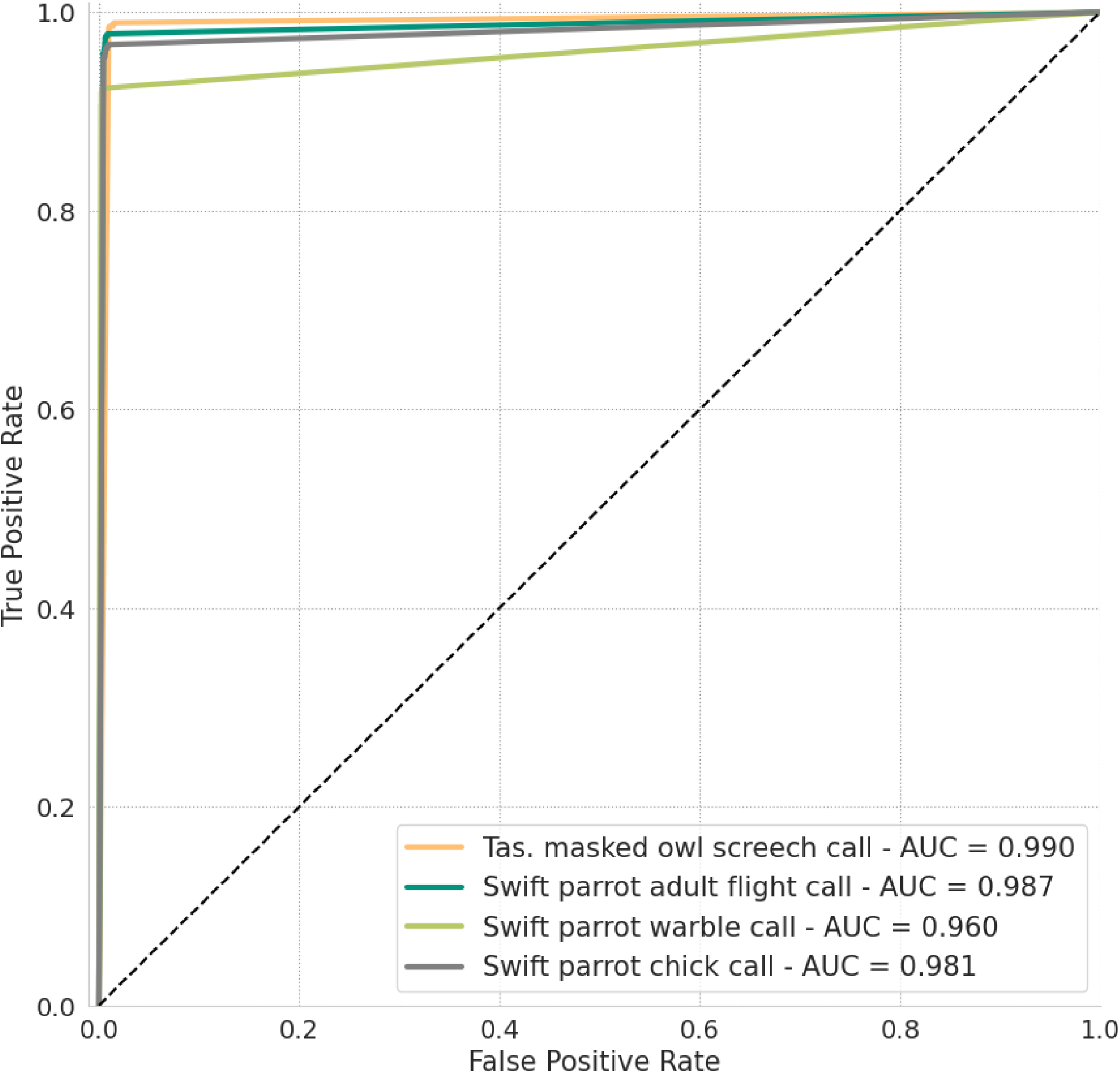

## Appendix D: Tasmanian masked owl survey results at the TI013E area

| **Site** | **Habitat type** | **Number of calls** | **Nightly detection rate (%)** |
| --- | --- | --- | --- |
| 1 | Mature Eucalyptus forest, impacted by recent wildfire | 19 | 35.7 |
| 2 | Mature Eucalyptus forest, impacted by recent wildfire | 3 | 10.5 |
| 3 | Young Eucalyptus forest, after clearfelling logging | 51 | 52.2 |
| 4 | Mature Eucalyptus forest, impacted by recent wildfire | 42 | 55.6 |
| 5 | Mature Eucalyptus forest, impacted by recent wildfire | 22 | 78.6 |
| 6 | Mature Eucalyptus forest, impacted by recent wildfire | 11 | 15.8 |
| 7 | Young Eucalyptus forest, after clearfelling logging | 10 | 42.9 |
| 8 | Mature Eucalyptus forest, with rainforest understorey | 53 | 68.2 |
| 9 | Mature Eucalyptus forest, with rainforest understorey | 177 | 95.7 |
| 10 | Rainforest | 37 | 72.2 |
| 11 | Young Eucalyptus forest, after clearfelling logging | 51 | 35.0 |
| 12 | Mature Eucalyptus forest, with rainforest understorey | 36 | 56.5 |
| 13 | Mature Eucalyptus forest, with rainforest understorey | 153 | 78.3 |
| 14 | Young Eucalyptus forest, after clearfelling logging | 28 | 60.0 |
| 15 | Rainforest | 99 | 71.4 |
| 16 | Young Eucalyptus forest, after clearfelling logging | 2 | 10.5 |
| 17 | Young Eucalyptus forest, after clearfelling logging | 8 | 22.2 |
| 18 | Young Eucalyptus forest, after clearfelling logging | 31 | 55.0 |

